# Trajectories of change after a health-education program in Japan

**DOI:** 10.1101/568584

**Authors:** MJ Park, Joseph Green, Hun Sik Jung, Yoon Soo Park

**Author notes:** Corresponding author: Joseph Green^3^, < >.

## Abstract

**Background:** Health education can benefit people with chronic diseases. However, in previous research those benefits were small, and reinforcement to maintain them was not effective. A possible explanation is that the benefits *appeared* to be small and reinforcement *appeared* to be ineffective because those analyses mixed data from two latent groups: one group of people who needed reinforcement and one group of people who did not. The hypothesis is that mixing the data from those two different groups caused the true effects to be “diluted.”

**Methods:** To test that hypothesis we used data from the Chronic Disease Self-Management Program in Japan, focusing on anxiety, depression, and patient-physician communication. To identify latent trajectories of change after the program, we used growth-mixture modeling. Then, to find out which baseline factors were associated with trajectory-group membership, we used logistic regression.

**Results:** Growth-mixture modeling revealed two trajectories – two groups that were defined by distinct patterns of change after the program. One of those patterns was improvement followed by backsliding: decay of impact. On anxiety and depression the decay of impact was large enough to be clinically important, and its prevalence was as high as 50%. Next, logistic regression analysis revealed that being in the decay-of-impact group could be predicted from multimorbidity, low self-efficacy, and high scores on anxiety or depression at baseline. In addition, one unexpected finding was an association between multimorbidity and *better* patient-physician communication.

**Conclusions:** These results support the hypothesis that previous findings (i.e. *apparently* small effect sizes and *apparently* ineffective reinforcement) actually reflect “dilution” of large effects, which was caused by mixing of data from distinct groups. Specifically, there was one group with decay of impact and one without. Thus, evaluations of health education should include analyses of trajectory-defined groups. These results show how the group of people who are most likely to need reinforcement can be identified even before the educational program begins. Extra attention and reinforcement can then be tailored. They can be focused specifically to benefit the people with the greatest need.

## INTRODUCTION

Nothing lasts forever. Yet a worthy goal of health education is for its benefits to be sustained. In some studies, benefits of health education have been found to last longer than 6 months (Barlow, Wright, Turner, & Bancroft, 2005; Barlow et al., 2008; Brady et al., 2013), while in others the findings are more nuanced – some improvements do not endure (Caplin & Creer, 2001; Clark, 2003; Franks, Chapman, Duberstein, & Jerant, 2009; Hennessy et al., 1999; Krebs, Prochaska, & Rossi, 2010; Lorig, Ritter, Laurent, & Fries, 2004; Norris, Lau, Smith, Schmid, & Engelgau, 2002). To sustain improvements, reinforcement has been recommended (Clark, 2003; Green, 1977; Newman, Steed, & Mulligan, 2009), but reinforcement has generally not been found to be useful (Glasgow, Toobert, Hampson, & Strycker, 2002; Lorig & Holman, 1989; Lorig, Ritter, Laurent, & Plant, 2006; Lorig, Ritter, Villa, & Piette, 2008; Nguyen, Carrieri-Kohlman, Rankin, Slaughter, & Stulbarg, 2005; Riemsma, Taal, & Rasker, 2003).

In this context, we consider the Chronic Disease Self-Management Program (CDSMP), which can improve health status and can increase the frequency of desirable health-related behaviors (Franek, 2013; Whitelaw, Lorig, Smith, & Ory, 2013). While the benefits of the CDSMP are statistically significant, some of them have also been described as minimal (Franek, 2013) or moderate (Brady et al., 2013). The program’s developers attributed these “modest” effect sizes to heterogeneity in the clinical and demographic characteristics of the participants (Lorig, Ritter, Laurent, & Plant, 2006), which points to a need to understand differences among participants and factors that might contribute to larger and more-sustained benefits.

In general, treating participants as homogeneous conceals true heterogeneity. For example, an intervention may be very useful in some participants, but that fact will not be recognized if one examines only the average for the group as a whole (Moynihan, Henry, & Moons, 2014). In addition, important heterogeneity in treatment effects can occur not only across groups but also over time. Sustained improvement in some participants could obscure relapse in others. Reinforcement given to all can appear to be ineffective, even if it is quite useful to some.

In previous research, subsets of participants were defined by socio-demographic characteristics, personality factors, etc., and not by patterns of change after the intervention (Franks et al, 2009; Harrison et al., 2012; Jerant, Chapman, Duberstein, & Franks, 2010; Reeves et al., 2008; Swerissen et al., 2006; Smeulders et al., 2010). LW Green (1977) discussed five distinct patterns of change after health education, and referred to the pattern in which good outcomes do not endure as *decay of impact* (also sometimes called backsliding). Since then, decay of impact as a person’s pattern of change after chronic-disease self-management education has almost never been studied. But knowing the magnitude of decay, and knowing when and in whom it occurs would be very useful in evaluating a health-education program’s effectiveness, and also in targeting interventions. Information about decay of impact can give planners an objective basis for deciding whether reinforcement is needed, when it is needed (Hennessy et al., 1999), and who is most likely to need it (Park, Green, Ishikawa, & Kiuchi, 2012).

Taking seriously the possibility of decay of impact entails defining groups by the trajectories of their outcomes during follow-up. Using data collected in two waves, Nolte, Elsworth, Sinclair, & Osborne (2007) may have been the first to study groups defined by their change after health education. An extension from patterns defined using two waves of data to those defined using four was published a few years later (Park et al., 2013). However, without the use of a well-established method for analyzing longitudinal data, doubts remain regarding different trajectories in outcomes. In this current study, using data collected before and after the CDSMP we applied Growth-Mixture Models (GMM) to empirically identify latent groups that were characterized by their patterns of longitudinal change, and we subsequently used logistic regression to identify baseline factors contributing to group membership (Cook, Karriker-Jaffe, Bond, & Lui, 2015; Hibbard, Mahoney, Stock, & Tusler, 2007; Muthén, 2004; Rabe-Hesketh, Skrondal, & Pickles, 2004; Ram & Grimm, 2009). This approach allows more granularity in evaluating the program’s effects and more accuracy in assessing needs for reinforcement.

## METHODS

### Participants and the program

Data were collected from participants in the CDSMP in Japan. They were recruited through public service centers and the Internet homepage of the Japan Chronic Disease Self-Management Association (Japan Chronic Disease Self-Management Association, 2018).

Based on self-efficacy theory, the program aims to build the participants’ skills in six areas: 1) handling pain, fatigue, frustration, and isolation, 2) exercising to maintain and increase strength, endurance, and flexibility, 3) using medications appropriately, 4) improving communication with friends, family, and healthcare professionals, 5) achieving and maintaining proper nutrition, and, 6) evaluating new therapies (Chronic Disease Self-Management Program, 2018). Skills in those areas were taught and practiced during group-discussion sessions that were held once each week for six weeks. Each group had two lay facilitators who had undergone approximately 35 hours of training. A textbook was used as the reference for the program’s content (Lorig, Holman, Sobel, Laurent, & González, 2001).

### Measures

Demographic and clinical information were collected using self-administered questionnaires. Also included in the questionnaires were scales to measure health status, health-related behaviors, psychological variables, etc. Those included self-efficacy to manage chronic health conditions (on a 0-to-60 scale) and the Hospital Anxiety and Depression Scale (HADS, Matsudaira et al., 2009). The HADS asks about symptoms of anxiety and of depression in the past week. Possible total scores on both the anxiety scale and the depression scale range from 0 to 21, with higher scores reflecting more symptoms and more frequent symptoms. Also included was a 3-item scale to measure communication with physicians, with possible total scores ranging from 0 to 15. Higher scores reflected more frequent use of proactive methods for good patient-physician communication (Lorig et al., 1996).

### Study design and timing of measurements

Data were collected four times over one year. Baseline data were collected before the first group-discussion session. Follow-up questionnaires were sent by postal mail 3, 6, and 12 months later. A post-paid self-addressed envelope was included. A reminder postcard was sent to each participant whose follow-up questionnaire was not received within two weeks.

### Analyses

To allow detection of decay of impact, the analyses were done using data from participants who provided at least three waves of data (456/643; 71%). Unconditional quadratic growth curves (Rabe-Hesketh, Skrondal, & Pickles, 2004) and conditional growth curves were fit using the four-wave data with all participants. Time-variant (self-efficacy) and time-invariant covariates (gender, educational status, partnered status, number of diagnoses, and history of illness) were included in the growth-curve analysis, with anxiety, depression, and communication as outcomes, to examine baseline differences and interactions over time. Quadratic terms were specified on “time” by including a squared time (i.e. time x time) term. Interaction terms between time-invariant covariates and time were included to examine factors contributing to longitudinal changes.

GMM with quadratic growth curves were fit for 2, 3, and 4 latent groups. We decided on the final number of groups after examining the relative fit (Bayesian Information Criterion [BIC]) and absolute fit (Proportion Correctly Classified ^[*P*^_*c*_] based on posterior probability). Multiple logistic regression was used to identify factors contributing to trajectory-group membership. Of the 456 participants whose data were analyzed by GMM here, 369 were included in a previous non-GMM analysis (Park et al., 2013). Data were analyzed with Latent Gold 5.1 (Belmont, MA), Stata 14 (College Station, TX), and JASP (https://jasp-stats.org/).

### Ethics

This study was approved by the University of Tokyo (number 1472–(2), Research Ethics Committee, Graduate School of Medicine). Participation in the CDSMP and in this research were voluntary. Informed consent was obtained in writing from all participants before the study began.

## RESULTS

### 1. The participants

Data from 456 participants were analyzed. Among them, 79% were women, 48% were college educated, 52% were partnered (married or living with someone), and 47% had more than one chronic condition. Details of multimorbidity are in Appendix 1.

### 2. All participants considered together

In the analysis with all participants considered together (Table 1), for the first six months communication with physicians increased, while both anxiety and depression decreased. However, by the end of the follow-up year all three outcomes had begun changing back toward their baseline values. That is, the initial improvements appeared to be followed by at least some backsliding.

**Table 1.**
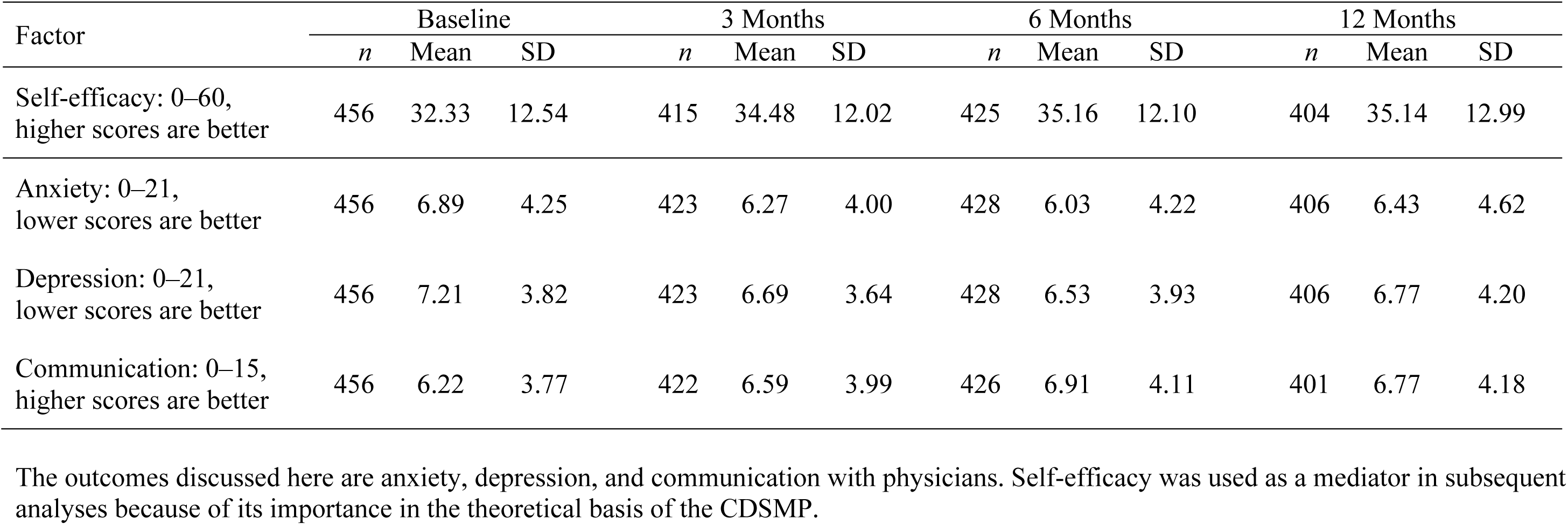
Descriptive statistics for all CDSMP participants considered together, baseline and follow up over one year.

Growth-curve analysis indicated that for all three outcomes the quadratic terms were significant (see Table 2). In addition, higher self-efficacy at baseline was associated with less anxiety, less depression, and better communication with physicians.

**Table 2.**
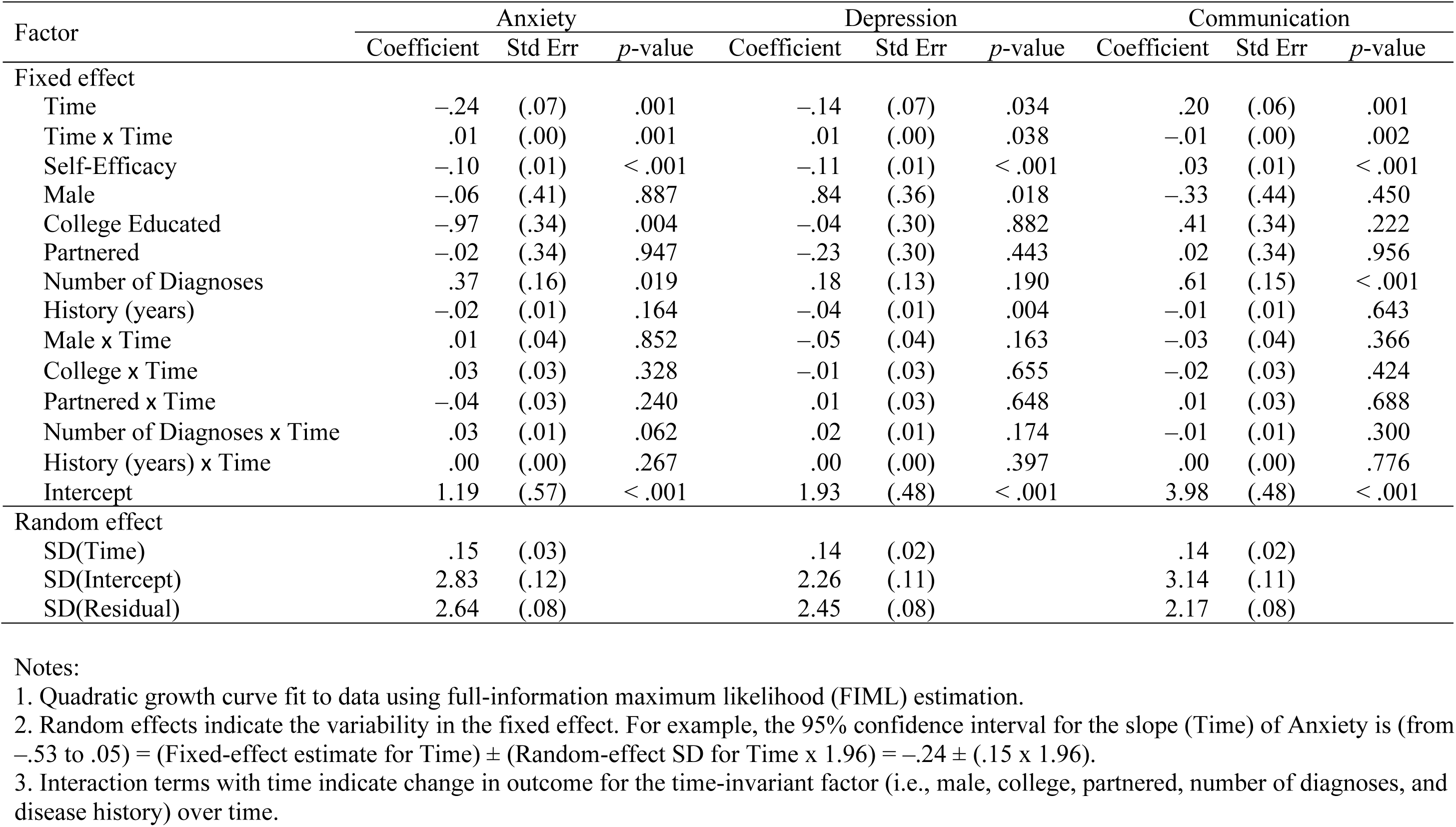
Results of growth-curve analysis, all CDSMP participants considered together (n = 456)

Also associated with both anxiety and communication was the number of diagnoses, and the regression coefficients for both were positive. That is, the participants with more comorbid conditions had greater anxiety at baseline. It is noteworthy that the participants with more comorbid conditions also had *better* baseline scores on the scale measuring communication with physicians. These associations did not change over time, as evidenced by the non-significant interaction term with time.

### 3. Groups defined by their trajectories

For all three outcomes, the GMM results were similar: The BIC and the *P*_*c*_ both led to the conclusion that the best-fitting models were those with two groups (Appendix 2).

For each outcome, those two groups began from substantially different baseline scores (Figure 1). Also for each outcome, one group changed very little throughout the follow-up period while the other changed noticeably within the first six months of follow-up, and then it reversed course back toward the baseline value (Figure 1). That is, on each outcome some participants were in a decay-of-impact group and the others were not. About half of the participants were in the decay-of-impact group: anxiety 45.6%, depression 50.7%, communication with physicians 46.3%.

**Figure 1.**
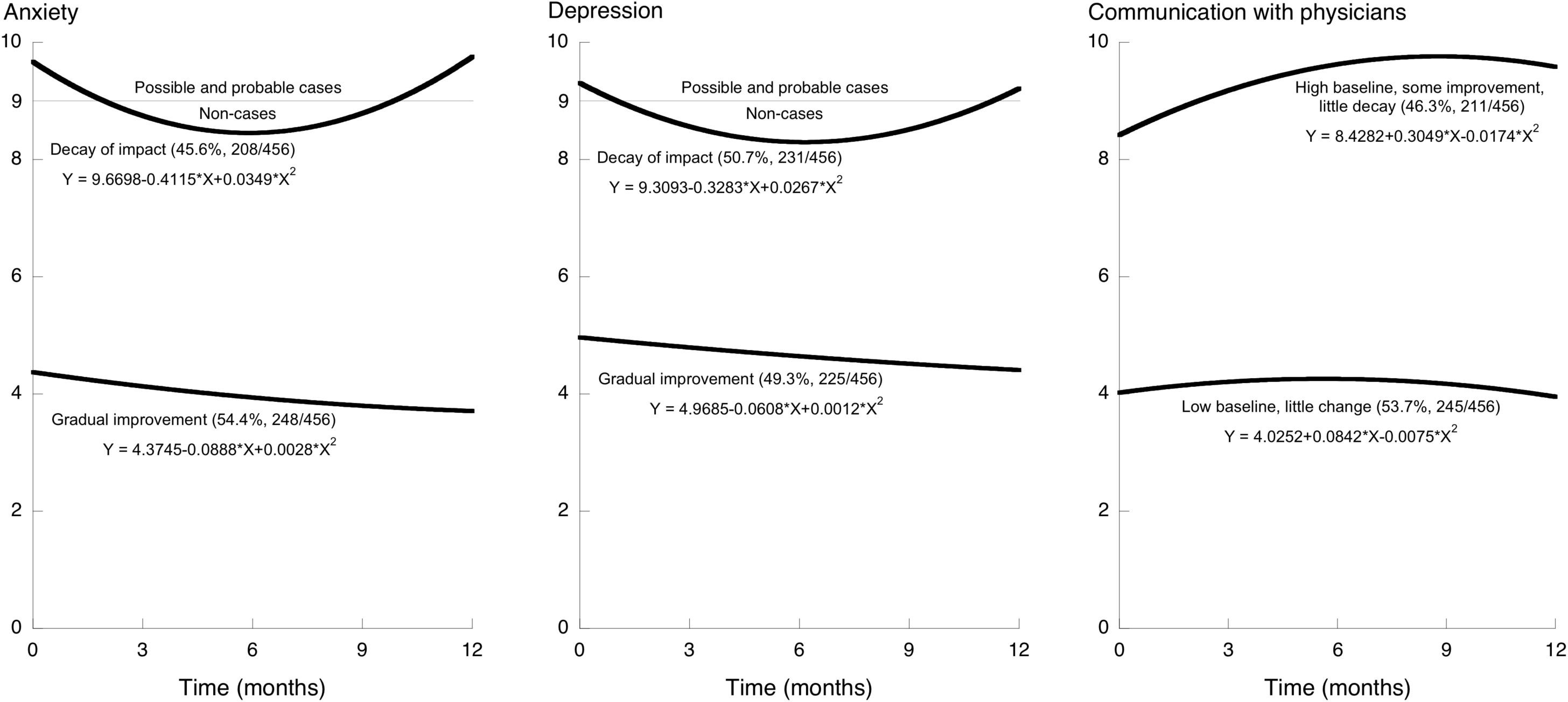
Trajectories of change after health education, showing two trajectory-defined groups for each of the three outcomes Growth-Mixture Modeling revealed two trajectory-defined groups for each outcome. On anxiety and depression higher scores are worse. On communication with physicians higher scores are better. For each outcome, one of those two groups had improvement followed by deterioration: decay of impact. For Anxiety and Depression, a score of 9 is the cutoff used in Japan to separate non-cases from possible and probable cases.

Participants who had decay of impact on one of the two mental-health outcomes (anxiety or depression) were also likely to be classified as having decay of impact on the other one (Phi = 0.508, Appendix 3). However, participants who had decay of impact on one of the two mental-health outcomes were no more or less likely to have decay of impact on communication (Phi = 0.095 and 0.043, Appendix 3).

On the two mental-health outcomes, the decay-of-impact group was the group with worse baseline status: more symptoms, and more-frequent symptoms, of anxiety and depression. In contrast, on communication with physicians the decay-of-impact group was the group with better baseline status: more frequent use of the three specified methods for good patient-physician communication. Also, by the end of the follow-up year the anxiety and depression scores had decayed back to their respective baseline levels, whereas on communication the decay trajectory was clear but the scores did not return to the baseline level, in other words the decay itself was smaller on communication with physicians than on the mental-health outcomes (Figure 1).

### 4. Contributors to group membership (Table 3)

For all three outcomes, self-efficacy at baseline was associated with group membership. Participants with higher self-efficacy were more likely to be in the group with lower anxiety at baseline, in the group with lower depression at baseline, and in the group with better communication at baseline.

**Table 3.**
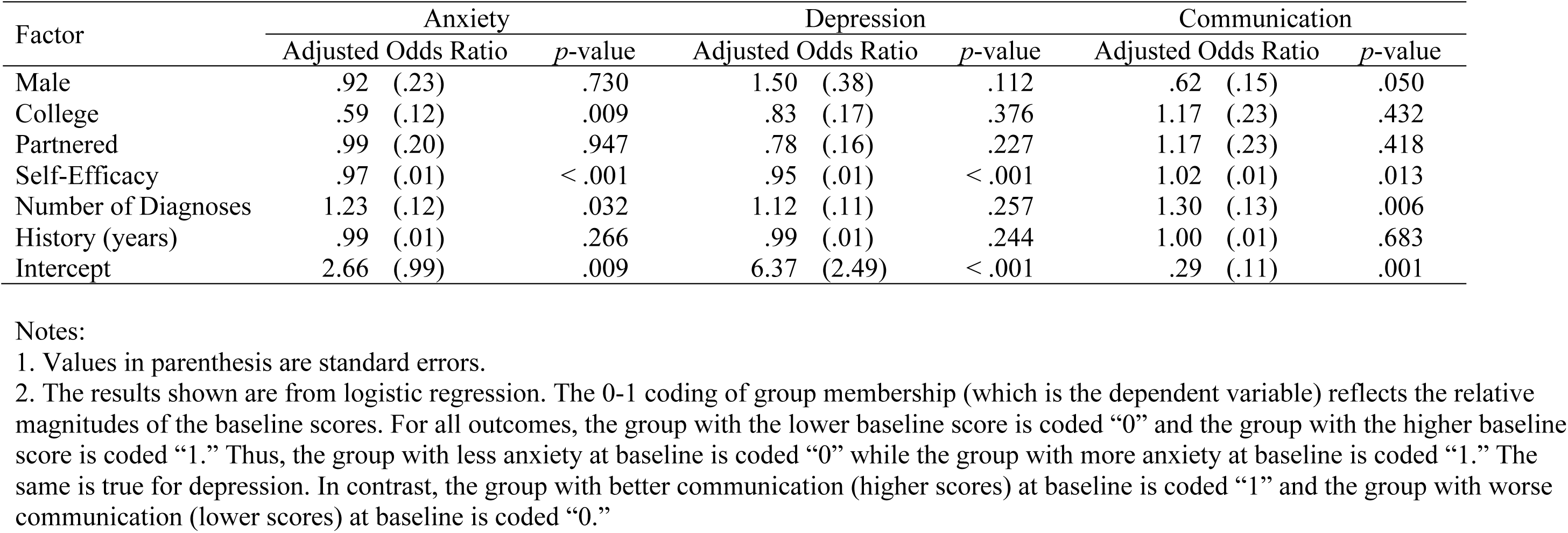
Factors predicting membership in groups defined by their trajectory after the CDSMP.

| Factor | Anxiety |  |  | Depression |  |  | Communication |  |  |
| --- | --- | --- | --- | --- | --- | --- | --- | --- | --- |
|  | Adjusted Odds Ratio |  | *p*-value | Adjusted Odds Ratio |  | *p*-value | Adjusted Odds Ratio |  | *p*-value |
| Male | .92 | (.23) | .730 | 1.50 | (.38) | .112 | .62 | (.15) | .050 |
| College | .59 | (.12) | .009 | .83 | (.17) | .376 | 1.17 | (.23) | .432 |
| Partnered | .99 | (.20) | .947 | .78 | (.16) | .227 | 1.17 | (.23) | .418 |
| Self-Efficacy | .97 | (.01) | < .001 | .95 | (.01) | < .001 | 1.02 | (.01) | .013 |
| Number of Diagnoses | 1.23 | (.12) | .032 | 1.12 | (.11) | .257 | 1.30 | (.13) | .006 |
| History (years) | .99 | (.01) | .266 | .99 | (.01) | .244 | 1.00 | (.01) | .683 |
| Intercept | 2.66 | (.99) | .009 | 6.37 | (2.49) | < .001 | .29 | (.11) | .001 |

Notes:
1. Values in parenthesis are standard errors. 2. The results shown are from logistic regression. The 0-1 coding of group membership (which is the dependent variable) reflects the relative magnitudes of the baseline scores. For all outcomes, the group with the lower baseline score is coded “0” and the group with the higher baseline score is coded “1.” Thus, the group with less anxiety at baseline is coded “0” while the group with more anxiety at baseline is coded “1.” The same is true for depression. In contrast, the group with better communication (higher scores) at baseline is coded “1” and the group with worse communication (lower scores) at baseline is coded “0.”

For anxiety and for communication with physicians, the number of diagnoses was also associated with group membership. Regarding anxiety, participants with more diagnoses were more likely to be in the group with higher (i.e. worse) scores at baseline and subsequent decay of impact. Regarding communication, participants with more diagnoses were more likely to be in the group with higher (i.e. *better*) scores at baseline and subsequent decay of impact.

## DISCUSSION

### All participants

When all participants were considered together, all three outcomes improved over the first six months. That improvement was followed by a small deterioration. Thus, even from the least-detailed analyses, some decay of impact was evident (Table 1). That interpretation is supported by the results of the growth-curve analyses: the quadratic terms were significant for all three outcomes (Table 2).

The growth-curve analyses also showed that higher self-efficacy at baseline was associated with less anxiety, less depression, and better communication with physicians, which is consistent with the theoretical basis of the CDSMP (Lorig & González, 1992).

Also evident at this level of analysis were associations with multimorbidity. Having more diagnoses was associated with more anxiety, more depression, and *better* communication with physicians. Of those three findings, the first two might be expected, but the third is particularly interesting. It is also reflected in the group-membership analyses, and so we will discuss it below.

### Findings from GMM: groups defined by their trajectories

For all three outcomes the results of GMM were consistent: Among all of the models tested, the two-group models had the lowest BIC and the highest *P*_*c*_. We are therefore confident in saying that GMM revealed *two* latent groups among these participants in the CDSMP. In some circumstances practical considerations could override the conclusions from those statistical criteria, as described in Appendix 2.

As noted above in the Introduction, in some previous studies, subsets of CDSMP participants were defined *a priori* and with reference to theory. In contrast, groups identified by GMM are empirical, as is each participant’s group membership. We note that the GMM approach can lead to testable hypotheses (regarding multimorbidity, as described below), and it can be used to answer important questions about whether similar phenomena also occur among other groups and in different settings.

### Mental health, and reinforcement

For both mental-health outcomes, the trajectory-defined groups differed in their baseline status and in their pattern of change after the program. Regarding anxiety, approximately half of the participants were in a group that began with relatively good scores, and they improved very gradually over the following year. In contrast, the other half were in a group that began from a high-anxiety baseline. That second group improved over the first 6 months, but by the time of the 12-month follow-up it had returned to its baseline level, and thus we refer to the latter group as the decay-of-impact group. The same was true with regard to depression.

Dichotomization is undoubtedly dangerous (Harrell, 2019), and yet HADS scores are used to separate people into categories of anxiety and depression severity. In Japan, the HADS threshold score separating non-cases from possible and probable cases was 9 (Matsudaira et al., 2009). The decay-of-impact trajectories on both anxiety and depression crossed that threshold twice – first during the improvement occurring soon after the intervention ended, and then again about 8 months later during the decay back toward the baseline value (Figures 1a and 1b). Therefore, to the extent that the threshold of 9 is useful, both the improvement measured soon after the CDSMP and also the deterioration measured near the end of the follow-up year were clinically important.

“Average therapeutic trial results can mislead” (Moynihan et al., 2014), but GMM provides more detail than average results. Here GMM showed that only some of the participants had decay of impact. At least on anxiety and depression, both the existence of a decay-of-impact group and the movement of that group between clinical categories support the idea that follow-up interventions – reinforcement – should be offered to *some* of the participants. Had reinforcement been given to all, it is unlikely that those in the group without decay of impact would have benefitted from it, simply because they already had almost no psychological distress – almost no room to improve. Rather than being expended on all of the participants, the resources used to implement reinforcement should be saved for the people who need it, to help them maintain their newly-improved status or perhaps improve further.

The present findings show how the small effect sizes and null results in published studies of reinforcement could be underestimates. Specifically, if all participants are considered together then any benefits to the decay-of-impact group will be diluted. Testing the effects of reinforcement is reasonable, but only in those whom reinforcement could benefit – the decay-of-impact group. To identify that group at baseline, i.e., even before the CDSMP begins, the present findings offer two potential criteria: a high baseline score (greater distress) on the mental-health outcome of interest, and a low level of self-efficacy. With regard to anxiety, a third criterion could be multimorbidity.

### Communication with physicians

Similar to the results described above for mental health, regarding communication with physicians GMM revealed two groups, each comprising about half of the participants, and those two groups began from noticeably different baselines. One group (n = 245) started from a very low baseline communication score (about 4 points) and it changed very little over the following year (Figure 1). This could well indicate an unmet need. Specifically, by the standard implied in the patient-physician communication scale, for those 245 participants substantial improvement after the baseline measurement was possible, but it did not occur. This leads to at least three research questions: (1) Are some participants in fact satisfied with a “low” level of communication? (2) Was the program implemented as well as possible? and (3) Even if the implementation was good, would those 245 participants have benefitted from a more-intensive intervention with an even-greater emphasis on practicing communication skills?

Also noteworthy are the three criteria that might be used to pre-emptively identify the participants who are most likely to need communication-skill practice: a low communication score at baseline, low self-efficacy, and *uni*morbidity.

### Multimorbidity

The participants with more diagnoses had better communication scores in the initial growth-curve analysis (Table 2), and they were more likely to be in the trajectory-defined group that had better communication scores throughout the year (Table 3). While the CDSMP has been found to be particularly useful to people with multiple diagnoses (Harrison et al., 2012), here multimorbidity was associated with a desirable health-related behavior even at baseline. To address the apparent connection between having multiple diagnoses and communicating well with physicians, we begin by noting that the communication scores reflect how often the respondents do the following three activities: making a list of questions to ask one’s physician during clinic visits; asking one’s physician about things that one wants to know or does not understand regarding one’s treatment; and discussing (with one’s physician) personal problems related to one’s medical condition. The people with multiple diagnoses probably had more experience being in health-related situations that were difficult to manage. To deal with those difficulties, perhaps they began writing lists of questions, asking for clarification, and discussing personal issues related to their diseases, simply because their health conditions were so complex. We hypothesize that at least some of the people with multimorbidity had become accustomed to doing those activities, and therefore in the domain of patient-physician communication they had already become “expert patients” (Reeves et al., 2008) by the time the study began. To the extent that better patient-physician communication results in better clinical care, this connection between multimorbidity and good communication could account at least in part for the documented association of multimorbidity with higher-quality care (Min et al., 2007).

Other factors could also be important. For example, self-selection might have played a role. After all, participation in the program was voluntary (as it is worldwide). Personality can moderate the effects of the CDSMP (Franks et al., 2009; Jerant et al., 2010), and perhaps it also affects one’s decision to participate. Among all of the eligible people with multimorbidity, those who are less “conscientious” and less interested in self-managing their conditions would not often make lists of questions, etc., and they might not have found the CDSMP to be attractive and thus would not have participated. In contrast, the CDSMP might appeal to people like the highly-proactive communicator with 8 chronic conditions who was described by Haslam (2015). People with multiple diagnoses who take initiative in self-managing their condition(s) by writing lists of questions, etc. could be over-represented among the program’s participants.

As explanations of the association between multimorbidity and good patient-physician communication, both the self-selection hypothesis and the already-an-expert-patient hypothesis remain to be tested, and of course both could be true.

### Limitations

The four waves of data collection over one year were more than enough to allow detection of decay of impact, but more frequent measurement and longer follow-up would of course be useful.

The number of diagnoses was self-reported. While we would have preferred to use medical records, for many chronic conditions self-reported diagnosis is accurate enough for research (Karison et al., 1999; Wada et al., 2009).

## Summary

GMM exposed two trajectory-defined groups, and the CDSMP clearly benefitted one group more than the other. However, the group that benefitted also had substantial decay of impact, and thus needed reinforcement. The decay-of-impact group comprised almost half of the participants. At baseline (i.e., before the program began), the participants most likely to need reinforcement were those with multimorbidity, those with low self-efficacy, and those who were clinically anxious or depressed.

Once the participants who are likely to have decay of impact are identified, extra attention and reinforcement can then be tailored. They can be focused specifically to benefit the people with the greatest need.

## Acknowledgments

The authors would like to express their gratitude to the Japan Chronic Disease Self-Management Association, as well as to all of the people who participated in the study. MJ Park is grateful for advice received from Y Yamazaki, for help and collaboration provided by the self-management research team at the University of Tokyo, for administrative assistance and technical support received from N Okamoto, and for advice and academic support received from T Kiuchi and H Ishikawa.

---

Appendix Table 1.

**Appendix Table 1a.** Participants in the study: basic descriptive statistics

| **Statistic** | **History of illness (years)** | **Age (years)** | **Number of Diagnoses** |
| --- | --- | --- | --- |
| **Valid n** | 441 | 453 | 456 |
| **Number Missing** | 15 | 3 | 0 |
| **Mean** | 13.64 | 48.79 | 1.774 |
| **Median** | 10.00 | 48.00 | 1.000 |
| **Std. Deviation** | 12.15 | 14.03 | 1.052 |
| **Skewness** | 1.313 | 0.05549 | 1.633 |
| **Std. Error of Skewness** | 0.1162 | 0.1147 | 0.1143 |
| **Kurtosis** | 1.559 | -0.7745 | 2.972 |
| **Std. Error of Kurtosis** | 0.2320 | 0.2289 | 0.2282 |
| **Minimum** | 0.000 | 18.00 | 1.000 |
| **25th percentile** | 4.000 | 38.00 | 1.000 |
| **50th percentile** | 10.00 | 48.00 | 1.000 |
| **75th percentile** | 20.00 | 59.00 | 2.000 |
| **Maximum** | 63.00 | 83.00 | 7.000 |

Number (%) of women: 362 (79%)
Number (%) living with spouse or partner: 238 (52%)

**Appendix Figure 1a.**
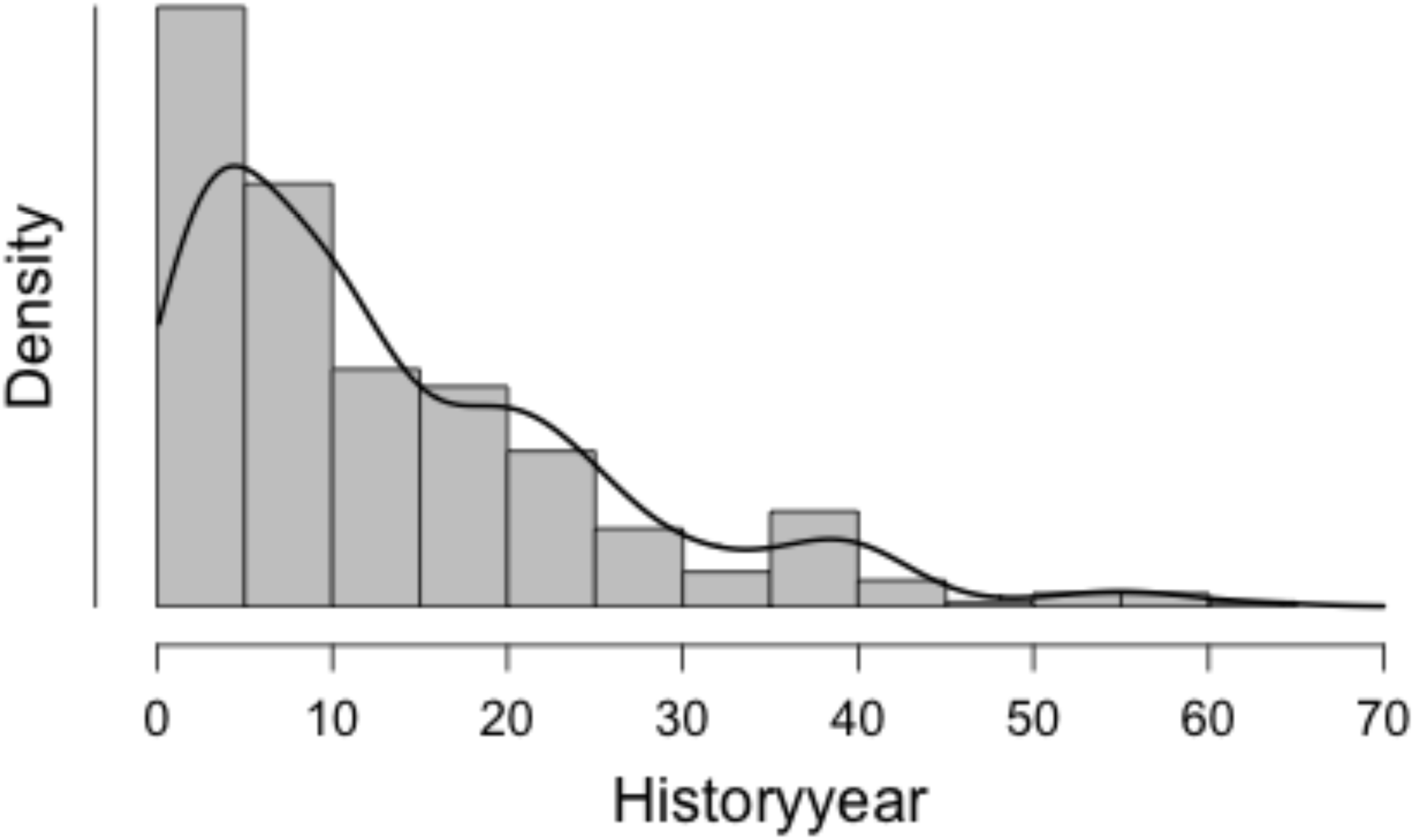
Frequency distribution of history of illness (years)

**Appendix Figure 1b.**
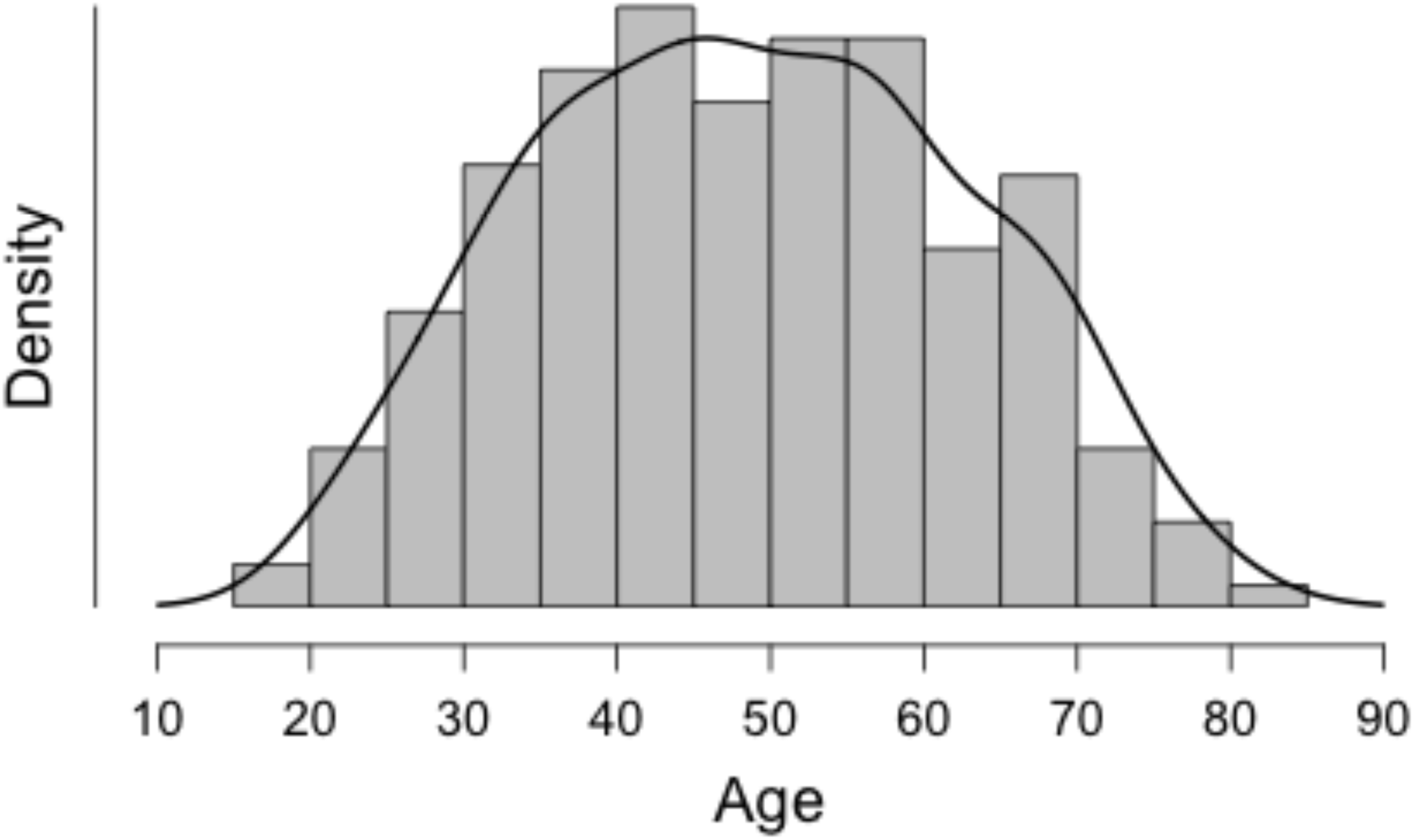
Frequency distribution of age (years)

**Appendix Figure 1c.**
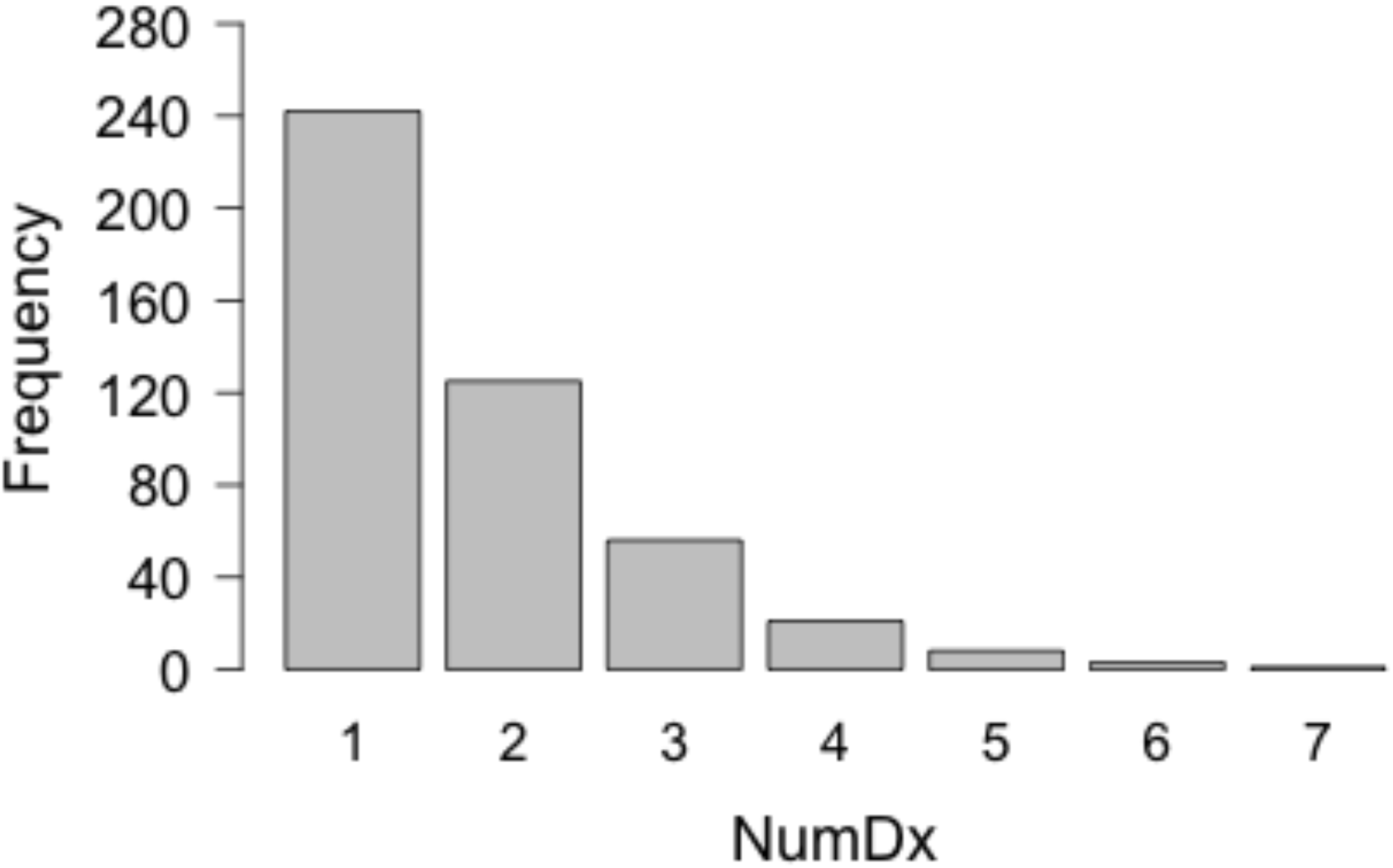
Frequency distribution of number of diagnoses

**Appendix Table 1b.**
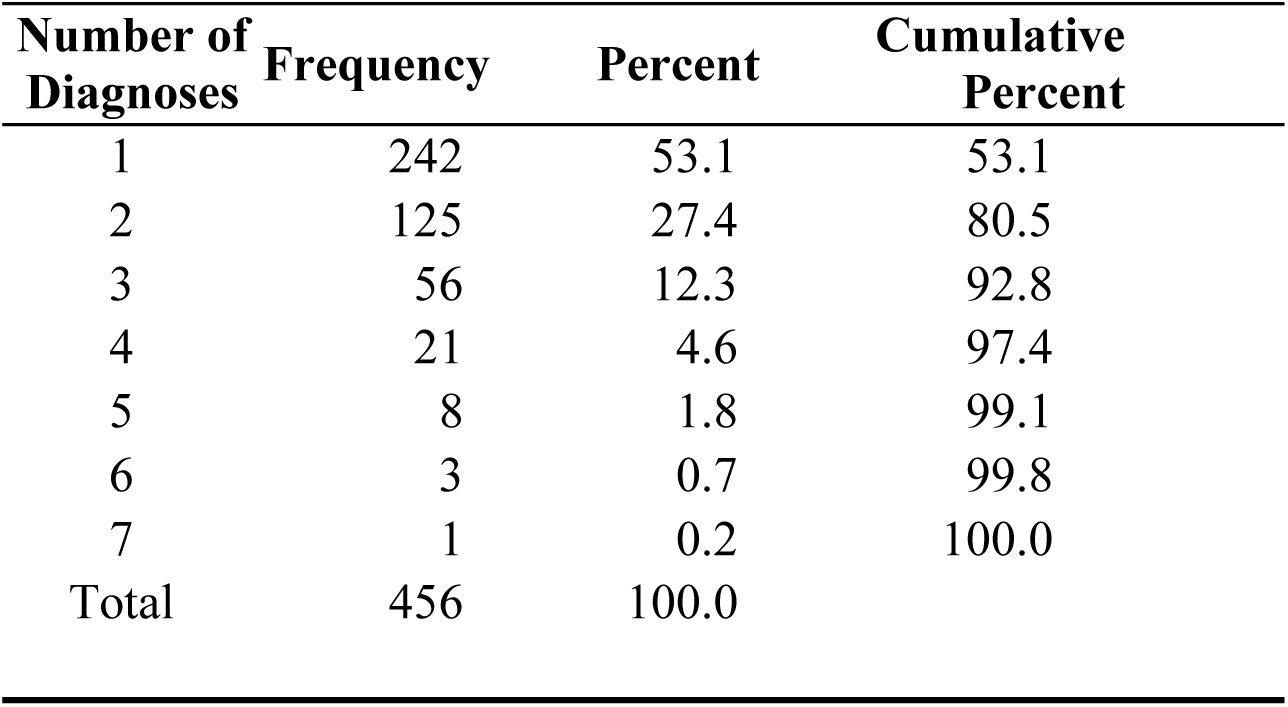
Frequencies of numbers of diagnoses

| Number of Diagnoses | Frequency | Percent | Cumulative Percent |
| --- | --- | --- | --- |
| 1 | 242 | 53.1 | 53.1 |
| 2 | 125 | 27.4 | 80.5 |
| 3 | 56 | 12.3 | 92.8 |
| 4 | 21 | 4.6 | 97.4 |
| 5 | 8 | 1.8 | 99.1 |
| 6 | 3 | 0.7 | 99.8 |
| 7 | 1 | 0.2 | 100.0 |
| Total | 456 | 100.0 |  |

**Appendix Table 1c.** Diagnoses, ordered by the number of participants with each diagnosis

| Diagnosis | Number of participants |
| --- | --- |
| Allergic disease | 120 |
| Cardiovascular disease | 96 |
| Connective tissue disease | 85 |
| Diabetes mellitus | 83 |
| Rheumatoid arthritis | 51 |
| Fibromyalgia syndrome | 38 |
| Cancer | 26 |
| Depression | 26 |
| Asthma | 23 |
| Inflammatory bowel disease | 21 |
| Pulmonary disease | 21 |
| Parkinson's disease | 13 |

Appendix 2.

In this study the decision about the number of groups was based on the BIC and the *P*_*c*_, but in some circumstances practical considerations could override the conclusions from those statistical criteria. For example, if only very limited resources are available to give reinforcement to the participants who are at risk of decay of impact, then a three-group model might be preferred. The reason is that the three-group model identified a group that begins from a much worse baseline value and is also more “volatile,” that is, it has both greater improvement and greater decay. That group is smaller – about one third rather than one half of the total. The limited resources could then be focused on that relatively small group with the greatest need. Because there might be such a practical reason for using three-group results, and for the readers’ general information, those results are given in the Tables here in Appendix 2.

This Appendix shows the results of Growth-Mixture Modeling.

1. BIC is the Bayesian Information Criterion, estimated using the deviance (–2 log likelihood). The model with the lowest BIC is preferred.

2. Classification (*P*_*c*_) is the proportion correctly classified, based on posterior probability. The model with the highest *P*_*c*_ is preferred [Clogg, C. C. (1995). Latent class models. In G. Arminger, C. C. Clogg, & M. E. Sobel (Eds.), Handbook of statistical modeling for the social and behavioral sciences (pp. 311-359). New York: Plenum Press].

3. Within each model that has more than one group, the equations are listed in order of increasing baseline value (Y intercept).

**Appendix Table 2a.**
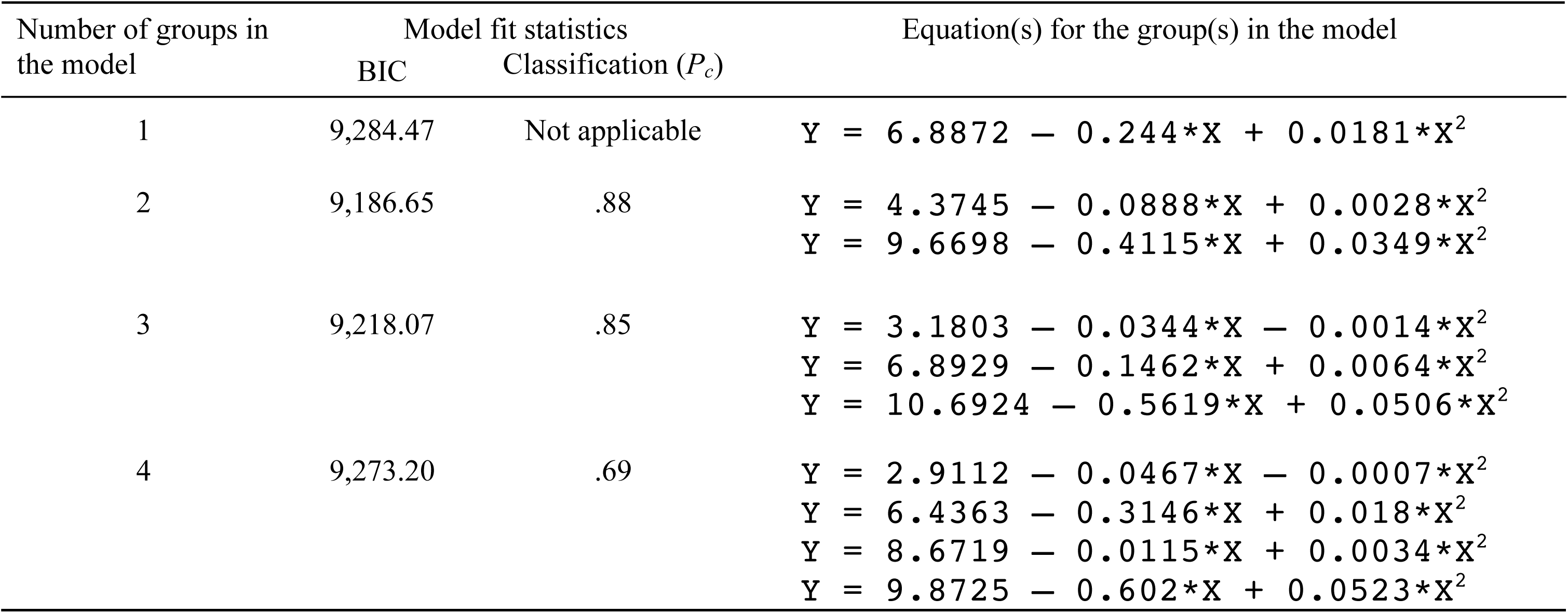
Four models of the Anxiety data

| Number of groups in<br>the model | Model fit statistics |  | Equation(s) for the group(s) in the model |
| --- | --- | --- | --- |
| | BIC | Classification ($P_c$) | |
| 1 | 9,284.47 | Not applicable | $Y = 6.8872 - 0.244 * X + 0.0181 * X^2$ |
| 2 | 9,186.65 | .88 | $Y = 4.3745 - 0.0888 * X + 0.0028 * X^2$<br>$Y = 9.6698 - 0.4115 * X + 0.0349 * X^2$ |
| 3 | 9,218.07 | .85 | $Y = 3.1803 - 0.0344 * X - 0.0014 * X^2$<br>$Y = 6.8929 - 0.1462 * X + 0.0064 * X^2$<br>$Y = 10.6924 - 0.5619 * X + 0.0506 * X^2$ |
| 4 | 9,273.20 | .69 | $Y = 2.9112 - 0.0467 * X - 0.0007 * X^2$<br>$Y = 6.4363 - 0.3146 * X + 0.018 * X^2$<br>$Y = 8.6719 - 0.0115 * X + 0.0034 * X^2$<br>$Y = 9.8725 - 0.602 * X + 0.0523 * X^2$ |

**Appendix Table 2b.** Four models of the Depression data

| Number of groups in<br>the model | Model fit statistics |  | Equation(s) for the group(s) in the model |
| --- | --- | --- | --- |
| | BIC | Classification ($P_c$) | |
| 1 | 9,022.85 | Not applicable | $Y = 7.1819 - 0.1966 * X + 0.0141 * X^2$ |
| 2 | 8,982.97 | .85 | $Y = 4.9685 - 0.0608 * x + 0.0012 * X^2$<br>$Y = 9.3093 - 0.3283 * x + 0.0267 * X^2$ |
| 3 | 9,018.32 | .80 | $Y = 4.1343 - 0.1223 * X + 0.0056 * X^2$<br>$Y = 6.7179 + 0.08 * X - 0.0047 * X^2$<br>$Y = 10.6954 - 0.5258 * X + 0.0395 * X^2$ |
| 4 | 9,082.24 | .65 | $Y = 4.3141 + 0.3974 * X - 0.0322 * X^2$<br>$Y = 5.9052 - 0.4869 * X + 0.0282 * X^2$<br>$Y = 6.9689 + 0.3035 * X - 0.0105 * X^2$<br>$Y = 11.8108 - 1.0304 * X + 0.0731 * X^2$ |

**Appendix Table 2c.** Four models of the Communication-with-physicians data

| Number of groups in<br>the model | Model fit statistics |  | Equation(s) for the group(s) in the model |
| --- | --- | --- | --- |
| | BIC | Classification ($P_c$) | |
| 1 | 8,690.05 | Not applicable | $Y = 6.2124 + 0.1951*X - 0.0124*X^2$ |
| 2 | 8,645.97 | .88 | $Y = 4.0252 + 0.0842*X - 0.0075*X^2$<br>$Y = 8.4282 + 0.3049*X - 0.0174*X^2$ |
| 3 | 8,671.51 | .72 | $Y = 3.4292 + 0.1005*X - 0.008*X^2$<br>$Y = 7.108 + 0.3629*X - 0.0197*X^2$<br>$Y = 8.4057 + 0.1401*X - 0.0108*X^2$ |
| 4 | 8,716.24 | .68 | $Y = 2.9075 + 0.1054*X - 0.0087*X^2$<br>$Y = 6.2069 - 0.0338*X - 0.0034*X^2$<br>$Y = 7.065 + 0.6366*X - 0.0289*X^2$<br>$Y = 8.7784 + 0.1061*X - 0.0103*X^2$ |

Appendix 3.

**Appendix Table 3.** Contingency tables for trajectory-group memberships (DoI: decay of impact)

a. Anxiety and depression
| Anxiety group | Depression group |  | Total |
| --- | --- | --- | --- |
|  | Not DoI | DoI |  |
| Not DoI | 180 | 68 | 248 |
| DoI | 45 | 163 | 208 |
| Total | 225 | 231 | 456 |

b. Anxiety and communication
| Anxiety group | Communication group |  | Total |
| --- | --- | --- | --- |
|  | Not DoI | DoI |  |
| Not DoI | 104 | 144 | 248 |
| DoI | 107 | 101 | 208 |
| Total | 211 | 245 | 456 |

c. Depression and communication
| Depression group | Communication group |  | Total |
| --- | --- | --- | --- |
|  | Not DoI | DoI |  |
| Not DoI | 109 | 116 | 225 |
| DoI | 102 | 129 | 231 |
| Total | 211 | 245 | 456 |

Phi coefficient = 0.043
95% CI: -0.053 to 0.138

